# Mining alternative splicing patterns in scRNA-seq data using scASfind

**DOI:** 10.1101/2023.08.19.553947

**Authors:** Yuyao Song, Guillermo Parada, Jimmy Tsz Hang Lee, Martin Hemberg

## Abstract

Single-cell RNA-seq is widely used for transcriptome profiling, but most analyses have focused on gene-level events, with much less attention devoted to alternative splicing. Here, we present scASfind, a novel computational method to allow for quantitative analysis of cell type-specific splicing events. scASfind utilizes an efficient data structure to store the percent spliced-in value for each splicing event. This makes it possible to exhaustively search for patterns among all differential splicing events, allowing us to identify marker events, mutually exclusive events, and large blocks of exons that are specific to one or more cell types. These methods allow researchers to compare cells based on isoforms rather than genes, thereby enabling more nuanced characterization of cell types and states. We demonstrate the advantages of scASfind on two mouse and one human datasets, identifying differences across the several key genes that cannot be detected using gene expression alone.

## Introduction

Alternative splicing (AS) is an essential, ubiquitous regulatory mechanism in eukaryotes. Through AS, a single gene can yield multiple mRNA isoforms, greatly expanding the protein diversity encoded by eukaryotic genes (Nilsen & Graveley, 2010). Fine-tuned regulation of alternative splicing events has a critical role in the development and function of diverse range of tissues and cell-types, including muscles, neurons, and immune cells (Baralle & Giudice, 2017; Dai et al., 2010; Ergun et al., 2013; Nikonova et al., 2020; X. Zhang et al., 2016). Splicing errors can also lead to an array of human diseases, such as neurodegenerative diseases, autoimmunity and cancer (Ren et al., 2021; Scotti & Swanson, 2015; Y. Zhang et al., 2021).

Decades of research using bulk methods have shown that many AS events are tissue-regulated, yet cell type-specific splicing remains incompletely understood. Using single cell RNA-seq (scRNA-seq), cell types can be comprehensively identified based on their expression profile, paving the way for studying splicing patterns. Although most scRNA-seq studies use droplet-based technologies such as 10X Chromium, which only profiles one end of the transcript, there are full-length scRNA-seq technologies, such as Smart-seq2 (Picelli, 2019) and VASA-seq (Salmen et al., 2022) that provide coverage of the entire transcript. Full-length technologies make it possible to conduct a local, event-level splicing quantification per cell type. In an event-level AS quantification, transcripts can be split into non-overlapping exonic regions, referred to as splicing nodes (Sterne-Weiler et al., 2018; Trapnell et al., 2010; Vaquero-Garcia et al., 2016). Nodes are further classified based on their behavior during splicing, e.g. core exons (CE) or alternative donors (AD). Then, the percent spliced-in (PSI) value for splicing nodes can be calculated based on reads spanning node junctions (Katz et al., 2010; Sterne-Weiler et al., 2018; Vaquero-Garcia et al., 2016). PSI is an informative indicator of exon usage frequency, providing an intuitive and easily interpretable metric to describe complex splicing events.

There are numerous computational methods for event-level splicing quantification in bulk RNA-seq, such as MISO (Katz et al., 2010), dSpliceType (Deng & Zhu, 2014), rMATS (Shen et al., 2014), MAJIQ (Vaquero-Garcia et al., 2016) and SUPPA2 (Trincado et al., 2018), but they are poorly suited due to the high sparsity and large size of scRNA-seq datasets. To overcome these issues, several methods aiming to detect and quantify AS in single-cell data have been developed. They include SingleSplice (Welch et al., 2016) which compares biological variation and technical noise in a population of single cells to find genes with isoform usage differences. Expedition (Song et al., 2017) is a suite of tools that can detect differences among the usage of splicing modalities. Huang and Sanguinetti have developed BRIE and BRIE2 (Huang & Sanguinetti, 2017, 2021), which uses Bayesian models for PSI estimation to overcome sparsity. SICILIAN (Dehghannasiri et al., 2021) assigns probabilities to called splice junctions to improve precision for their detection, and SpliZ (Olivieri et al., 2022) generalizes PSI to enhance splicing quantification at the single-cell level. A recent software tool is MARVEL (Wen et al., 2023), which integrates splicing and gene expression analyses. However, MARVEL analysis is limited to splicing events involving a single exon and it can only detect differential splicing between pairs of cell types. None of the methods presented to date can leverage event-level splicing quantification to comprehensively characterize cell type-specific splicing patterns, involving either single or multiple exons, without using a parametric model or imputing missing values.

To facilitate de novo detection of cell type-specific AS events, we developed scASfind, a flexible and intuitive method for mining complex AS patterns in large single-cell datasets. scASfind is an open-source R package which is freely available at https://github.com/hemberg-lab/scASfind. scASfind uses a similar strategy as our previous work scfind (Lee et al., 2021) to compress the cell pool-to-node differential PSI matrix into an index. This efficient data structure enables rapid access to cell type-specific splicing events, making it possible to use an exhaustive approach when carrying out pattern searches across the entire dataset. Importantly, scASfind does not involve any imputation or model fitting, instead cells are pooled to avoid the challenges presented by sparse coverage. Moreover, there is no restriction on the number of exons or the inclusion/exclusion events involved in the pattern of interest. Building on these fast searches, scASfind allows interactive searching of cell type-specificity of splicing patterns, such as differential splicing, mutually exclusive exons and coordinated splicing events. We applied scASfind to mouse primary visual cortex (Tasic et al., 2016) mouse embryonic development(Salmen et al., 2022), and human fetal liver (Ranzoni et al., 2021) to characterize cell type-specific splicing patterns.

## Results

### Data compression enables fast searching of splicing patterns

scASfind takes full-length scRNA-seq data, such as Smart-seq2 (Picelli et al., 2013), RamDA-seq (Hayashi et al., 2018), Smart-seq3 (Hagemann-Jensen et al., 2020), VASA-seq (Salmen et al., 2022) and FLASH-seq (Hahaut et al., 2022) as input for splicing quantification. Tag-based methods such as 10X Genomics Chromium are unsuitable since they only capture the transcript’s 3’ or 5’ end, and typically do not provide enough reads that span splice junctions. It is assumed that the data has been clustered and annotated so that each cell is assigned a cell type. Several cells of the same type are first combined into cell pools to provide sufficient reads for robust and accurate PSI quantification with Whippet (Sterne-Weiler et al., 2018) using the MicroExonator workflow (Parada et al., 2021).

After obtaining a splicing node x cell pool matrix of PSI values (**Fig 1a**), scASfind first centers each column of the matrix to obtain the deviation of PSI values from the dataset mean (default: |ΔPSI| > 0.2). This is to capture the biologically informative PSI variation across cells. The differential PSI matrix is further split into two to encode positive (spliced-in) and negative (spliced-out) PSI values. In both matrices, a non-zero value indicates that there is differential inclusion or exclusion of the splicing node in that particular cell pool. By ensuring that the matrices are sparse we can achieve a high compression rate and fast pattern matching, even for large datasets.

**Figure 1.**
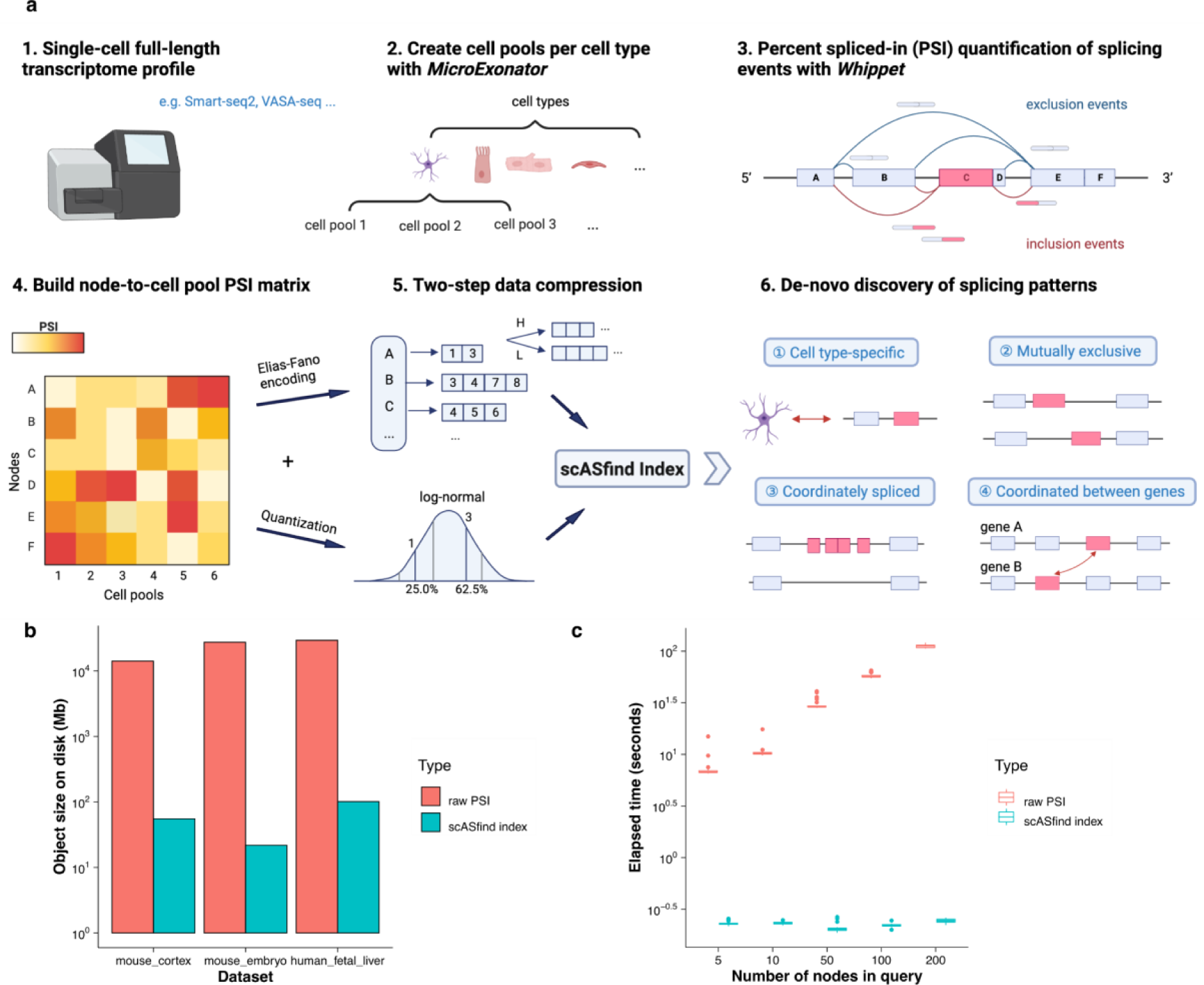
Overview of scASfind. (a) Schematic of scASfind workflow. Single-cell full-length transcriptome sequencing data such as Smart-seq2 or VASA-seq are suitable inputs for the scASfind workflow. Cells from the same cell type are pooled to increase the accuracy of splicing event detection (default 5 cells per pool) with MicroExonator (Parada et al., 2021). The PSI value for each splicing node is calculated by Whippet (Sterne-Weiler et al., 2018) to obtain a splice node-by-cell pool PSI matrix, and we then build a scASfind index containing information about splicing events that are differentially spliced in or spliced out in each cell pool. Finally, we query the index to search for cell type-specific differential splicing events, mutually exclusive node pairs and consecutive nodes that are similarly spliced-in or coordinated splicing events. (b) The size of the file saved to disk containing either the raw PSI values and metadata objects or the scASfind index with metadata objects built with a two bit quantization. (c) The elapsed time of searching all cells with increased inclusion, i.e. has a PSI no less than 0.2 higher than the dataset mean, in any of five randomly selected splicing nodes. The process is repeated 30 times. The bar in the boxplot shows the arithmetic mean, lower and upper hinges correspond to the first and third quartiles, whiskers extend from the hinge to the largest value no further than 1.5 * interquartile range from the hinge and outliers beyond this range are plotted as individual data points. PSI, percent spliced-in.

We adopted the indexing strategy in scfind (Lee et al., 2021) to compress the two sparse PSI matrices into two scASfind indexes, and we then combined them into a single meta-index object. The index efficiently stores the splicing nodes that have a PSI value deviated from the mean (see Methods for details about data compression into an index). In addition to providing efficient storage, the scASfind index also allows for rapid access to the raw PSI values associated with each splicing node. In particular, it allows us to use AND or OR queries to find the set of cell pools that match a set of inclusion/exclusion criteria, e.g. nodes 1, 2 and 3 need to be included well above mean while nodes 4 and 5 are excluded well below mean. By combining multiple queries we can carry out more complex searches, and since each operation is fast it becomes possible to adopt an exhaustive approach to search the entire dataset.

In the following, we set out to demonstrate how the scASfind index allows for thorough identification and characterization of cell type-specific splicing events. We used scASfind to analyze three technically and biologically distinct datasets: a mouse cortex data profiled using Smart-seq2 (hereafter referred to as mouse cortex) (Tasic et al., 2016), a mouse embryonic development dataset profiled using VASA-seq (hereafter referred to as mouse embryo) (Salmen et al., 2022), and a human fetal liver dataset profiled using Smart-seq2 (hereafter referred to as human fetal liver) (Ranzoni et al., 2021). For all three datasets, the scASfind representation required two to three orders of magnitude less disk space (**Fig 1B**) when using two bits for the quantizer. We also benchmarked the search times: compared to an implementation using only standard data structures, scASfind was hundreds or thousands of times faster (**Fig 1C**). For example, for the mouse embryo dataset, finding all pools that have increased inclusion of any of 200 randomly selected nodes with scASfind took 0.24 seconds on average, compared to 112 seconds for the naive approach. scASfind was also highly robust to increased search size. One additional overhead for scASfind is the time to build the index, but this is relatively minor as none of the datasets took more than 30 minutes (**Table 1**).

### Splicing events are more precise markers of cell types

Cell types are typically associated with a set of marker genes, i.e. genes that are highly expressed compared to other cell types. Since the most widely used single cell protocols do not provide enough information to distinguish isoforms, transcripts are usually evaluated at the gene level. However, identifying a reliable set of marker genes can be challenging, especially for neuronal tissues with complex cell type taxonomy (Kiselev et al., 2017). Since AS is known to be more prevalent in the brain (Dai et al., 2010; Hakim et al., 2017), we hypothesized that splicing events are more reliable for distinguishing cell types than gene expression. We refer to splicing nodes that are highly included or excluded in only one cell type as marker nodes in analogy with marker genes.

To identify cell type marker nodes, we used each cell type to query the scASfind index for nodes that have high or low inclusion. Benefiting from the speed at which these quantities can be extracted using the scASfind index, we carried out an exhaustive search to identify the best marker nodes for each cell type. Nodes were evaluated using the precision, recall and F1 score for their ability to detect the cell type of interest (see Methods for details). We used a similar procedure for marker genes using scfind, and we compared the quality of the markers by the precision, recall, and F1 scores. In the mouse cortex and the mouse embryo datasets, we observed higher F1 scores in splicing markers, compared with expression markers across the board (**Fig 2A, D**). Interestingly, the higher F1 of splicing markers is largely driven by higher precision (**Fig 2B, E**), suggesting that they yield few false positives. The F1 and precision of splicing and expression markers show comparative scores in the human fetal liver dataset (**Fig 2G, H**). The lack of benefit in using splicing markers for the human fetal liver suggests that the splicing landscape in this dataset is less complex compared to the mouse cortex and embryo, possibly due to the tissue. We conjecture that the poor recall (**Fig 2C, F, I**) for splicing markers could be due to the sparsity of splicing quantification leading to a high number of false negatives as there is insufficient information to accurately quantify splicing nodes in many pools.

**Figure 2.**
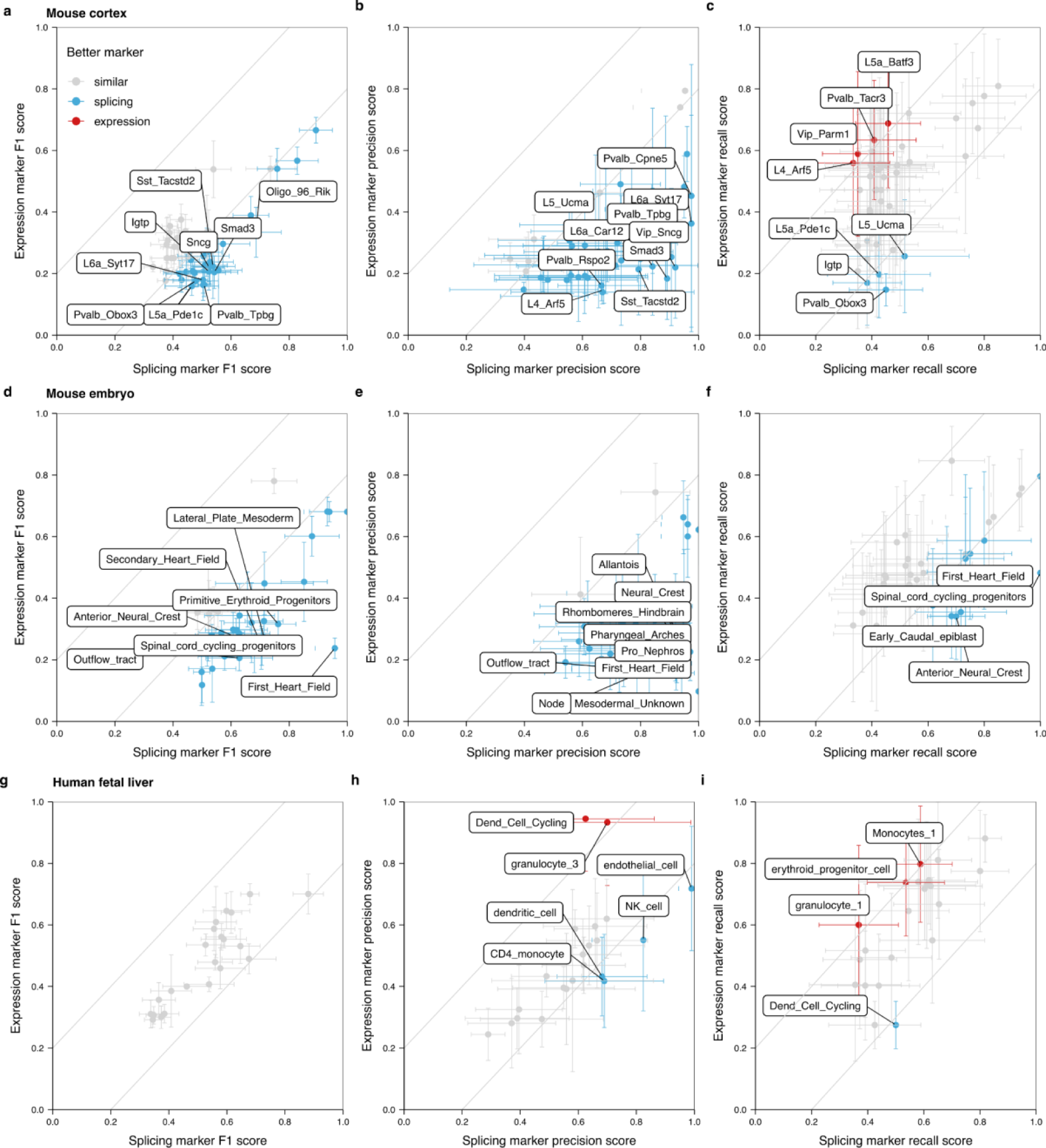
Comparing gene expression and splicing as cell type markers. The top 20 expression and splicing markers, ranked by F1 scores, and their precision, recall and F1 scores are calculated via scfind (Lee et al., 2021) or scASfind for (a-c) mouse cortex, (d-f) mouse embryo, and (g-i) human fetal liver. We compare the mean scores per cell type and consider a 0.2 difference between expression and splicing to indicate a better marker (gray lines in the figure). The dots represent the mean, while the whiskers indicate the minimum and maximum for the 20 markers. Cell types with either better splicing or expression markers are coloured (blue for splicing, red for expression). For visual clarity, cell types have score differences of 0.5, 0.6 or 0.2 for precision, 0.2, 0.3 or 0.2 for recall or 0.3, 0.4 or 0.2 for F1 in mouse cortex, mouse embryo and human fetal liver data are labeled, respectively. These values are chosen based on the respective number of cell types with a better marker for each dataset.

Moreover, the inclusion or exclusion of splicing nodes is independent of increased or decreased expression, suggesting that splicing markers are largely independent of the expression level (**Fig 3**). For instance, in astrocytes, *Dtna_27* is excluded while the cell type has higher expression of the *Dtna* gene. On the other hand, astrocytes have similar expression of the *Hnrnpa2b1* gene with other cell types while it has higher inclusion of *Hnrnpa2b1_32* (**Fig 3A, B**). Another example is glutamatergic neuron subtype L4_Scnn1a (**Fig 3C, D**), here we find both inclusion and exclusion splicing markers, while the expression levels of the corresponding genes can hardly distinguish this cell type from others. This is in line with observations by Wen et al. (Wen et al., 2023) that only a fraction of differentially spliced genes have expression changes in the same direction, indicating that differential splicing provides another layer of transcriptomics regulation that contributes to cell type heterogeneity.

**Figure 3.**
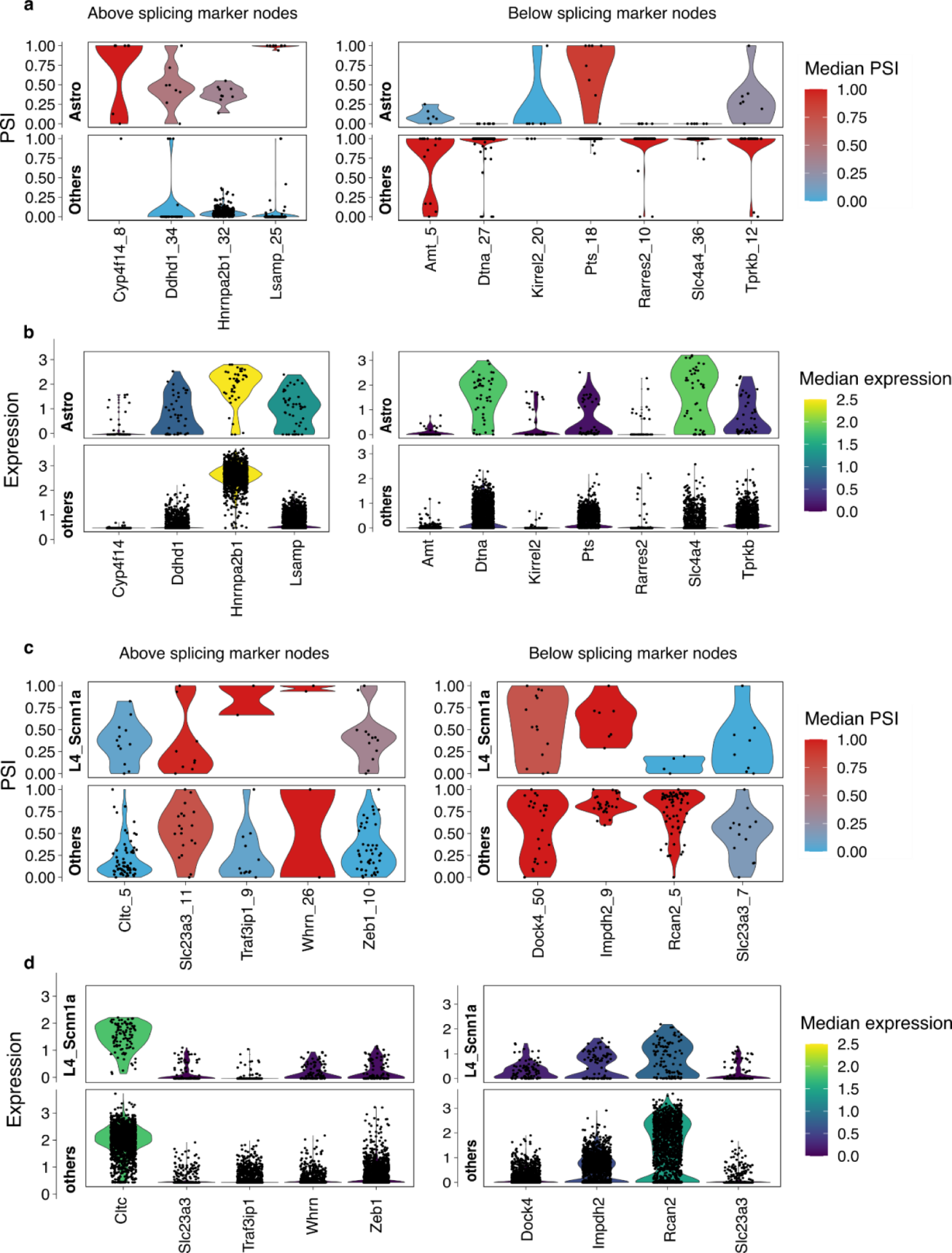
PSI values of marker splicing nodes and expression levels of corresponding genes. (a) PSI values for astrocyte splicing markers from mouse cortex data, compared to the mean of all other cell types. (b) Expression levels for the same genes contribute to splicing markers in astrocytes. (c) PSI values for L4_Scnn1a neuron splicing markers from mouse cortex data. (d) Expression levels for the same genes in (c). In all panels, each dot represents a cell pool and the color scale shows the median PSI or gene expression level among the cell pools. PSI, percent spliced-in.

While 67%, 57% and 62% of the top 20 splicing markers are from different genes in mouse cortex, mouse embryo and human fetal liver datasets, respectively, a single gene can contribute a large portion of marker nodes in some cases. We observed that the *Ttn* gene, which encodes Titin - the largest protein in the genome (Tharp et al., 2019), contributes most of the above splicing markers in the first heart field (**Fig 2F**) for the mouse embryo dataset. These results are consistent with previous analyses of Ttn splicing profiles that show 50-219 exons to be developmentally regulated (Baralle & Giudice, 2017). Taken together, this result suggests that splicing events frequently show higher cell-type specificity than gene expression, and the splicing marker events reported by scASfind can be superior in terms of distinguishing cell types. A particularly useful feature of splicing markers is their high precision, which is beneficial for rare cell types.

### Detecting mutually exclusive exon pairs

Mutually exclusive splicing events are a special type of AS event where only one of two consecutive exons is included in the final mRNA product (Hatje et al., 2017; Pohl et al., 2013). In the largest study of mutually exclusive exons (MXEs) in bulk RNAseq data carried out to date, Hatje et al identified 855 exon pairs from 515 datasets (Hakim et al., 2017; Hatje et al., 2017). MXE splicing is known to be regulated by different molecular mechanisms that enable tissue-specific patterns.(Kalsotra & Cooper, 2011; Pohl et al., 2013) (Rodriguez et al., 2020; Wang et al., 2008; Xu et al., 2002). We hypothesized that some of the observed tissue specific splicing profiles arise from the cumulative effects of cell type-specific MXE preferences within each tissue. Hence, we leveraged scASfind to systematically discover MXEs and explore their cell type specificity.

We performed an exhaustive search of all three datasets to observe cell type specificity for all exon pairs that could be MXEs (**Fig 4A**). That is, for all consecutive exons, we identified cell types in which one of them is always included and the other excluded. To ensure high-quality results, several additional filters were employed. First, we required the pair to have mean PSI values summing to 1±0.1, and that PSI standard deviation scores differ by less than 0.1 across all cell pools in the dataset. Second, we required at least one cell type to be significantly enriched for the pattern when one exon is included and the other is excluded. Statistical significance was determined using hypergeometric tests. Third, we considered MXEs detected in over half of the cell pools and having a difference of cell type mean PSI value between the two exons >= 0.5 as highly confident pairs. The default criteria we have used were rather stringent to obtain a small amount of highly confident results.

**Figure 4.**
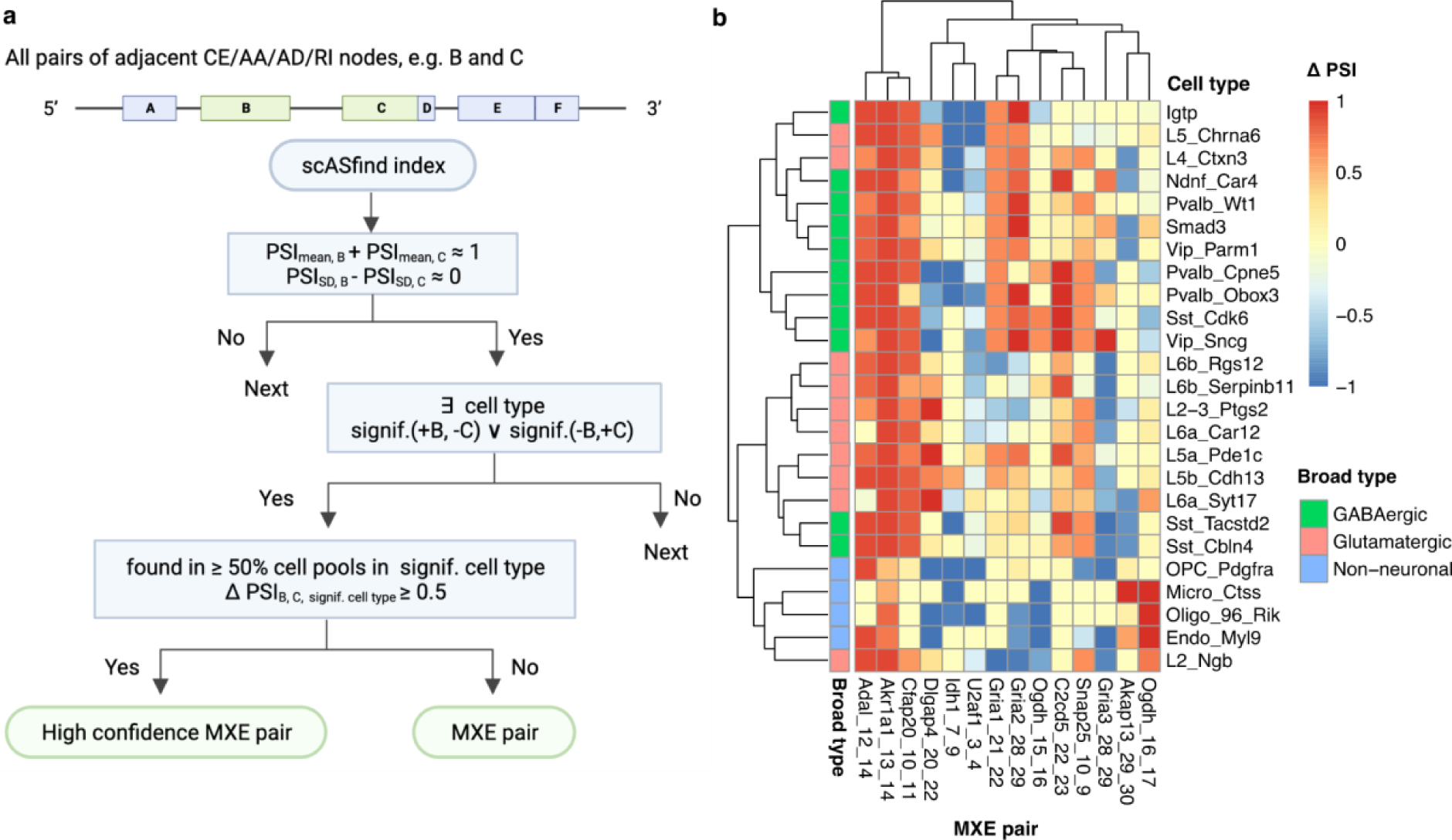
Summary of 14 high-confidence MXE pairs in mouse cortex data. (*a*) Schematic overview of how MXE pairs are identified. (*b*) High confidence adjacent mutually exclusive exon pairs detected by scASfind. The heatmap color scale indicates the difference in raw PSI value (ΔPSI) between the downstream exon and the upstream exon in the pairs of MXEs. Cell types are grouped by their broad type, genes are organized by the number of cell types in which the pair is significant (decrease from left to right). MXE, mutually exclusive exons; PSI, percent spliced-in; CE, core exon; AA, alternative acceptor; AD, alternative donor; RI, retained intron; signif., significant.

We detected 63, 17 and 35 significant pairs of MXEs in mouse cortex, mouse embryo and human fetal liver data, respectively. Among these 14, 2 and 2 were adjacent and highly confident. The high-confidence pairs in the mouse cortex dataset are summarized in **Fig 4B**. Hierarchical clustering across cell types using the ΔPSI of these exons indicates that generally, cells of the same broad type have similar splicing patterns in these MXEs, with a few exceptions. This is concordant with the tissue-specific splicing observed in bulk RNA-seq. However, there are examples of cell types within each broad type that have distinctive MXE preferences, suggesting a more complex pattern.

A known example of MXEs can be found in the ionotropic glutamate receptor genes, AMPA 1/2/3 (*Gria1*, *Gria2*, *Gria3*), which have been studied extensively in mouse brain (Koike et al., 2000; Sommer et al., 1990; Wright & Vissel, 2012). Reassuringly, the top candidates reported by scASfind include the three Gria genes. To the best of our knowledge, this is the first detailed study of cell-type-specific MXE preferences for these genes (**Fig 5A**). During development, some neurons switch from node 28 to using node 29, and this has important consequences for their responses to electric stimuli (Hadzic et al., 2017; Monyer et al., 1991). We detected a significant preference for node 29 in glutamatergic neurons including L2 Ngb and L2/3 Ptgst, as well as L6a Car12, L6b Rgs 12 and L6b_Serpinb11, GABA-ergic neurons including Vip_Sncg, Pvalb_Obox3, Smad3, Igtp and Pvalb_Wt1, as well as in oligodendrocytes Oligo_96_Rik. By contrast, several glutamatergic and GABA-ergic neurons included node 28, suggesting that the cell type-specific pattern is complex. Another example was in the SNARE protein *Snap25*, whose MXE preference switches during mouse brain development (Bark et al., 1995; Prescott & Chamberlain, 2011), and is related to regulating synaptic transmission and long-term synaptic plasticity (Boschert et al., 1996; Irfan et al., 2019). In our analysis, glutamatergic neurons L5_Chrna6, L6b_Rgs12, L6b_Serpinb11; GABA-ergic neuron Igtp, Ndnf_Car4, and oligodendrocyte progenitor cell OPC_Pdgfra showed a strong preference for node 9, while other cell types utilized node 10 (**Fig 5A**). Taken together, our results recapitulate some of the complex cell type specific pattern of isoform switching for both the Gria genes and *Snap25*. In the mouse embryo data, we detected highly confident MXEs in *Actn1* and *Actn4*. *Actn1* has been found to have tissue-specific mutually exclusive splicing in adult mice. Compared to other tissues, muscle cells select an alternative exon which makes the protein’s EF-hand motif insensitive to Ca2+, while brain cells include both exons (Kremerskothen et al., 2002; Waites et al., 1992). We were the first to describe the cell type preference of this MXE pair in the mouse embryo (**Fig 5B**). For *Actn4*, we found primitive_heart_tube cells prefer *Actn4_14* while first_heart_field and secondary_heart_field cells chose *Actn4_13.* Finally, for the *P4HA2* gene in the human fetal liver dataset, Common_Prog_cycling and Dend_cell_cycling selected node 47 or 48 respectively (**Fig 5C**).

**Figure 5.**
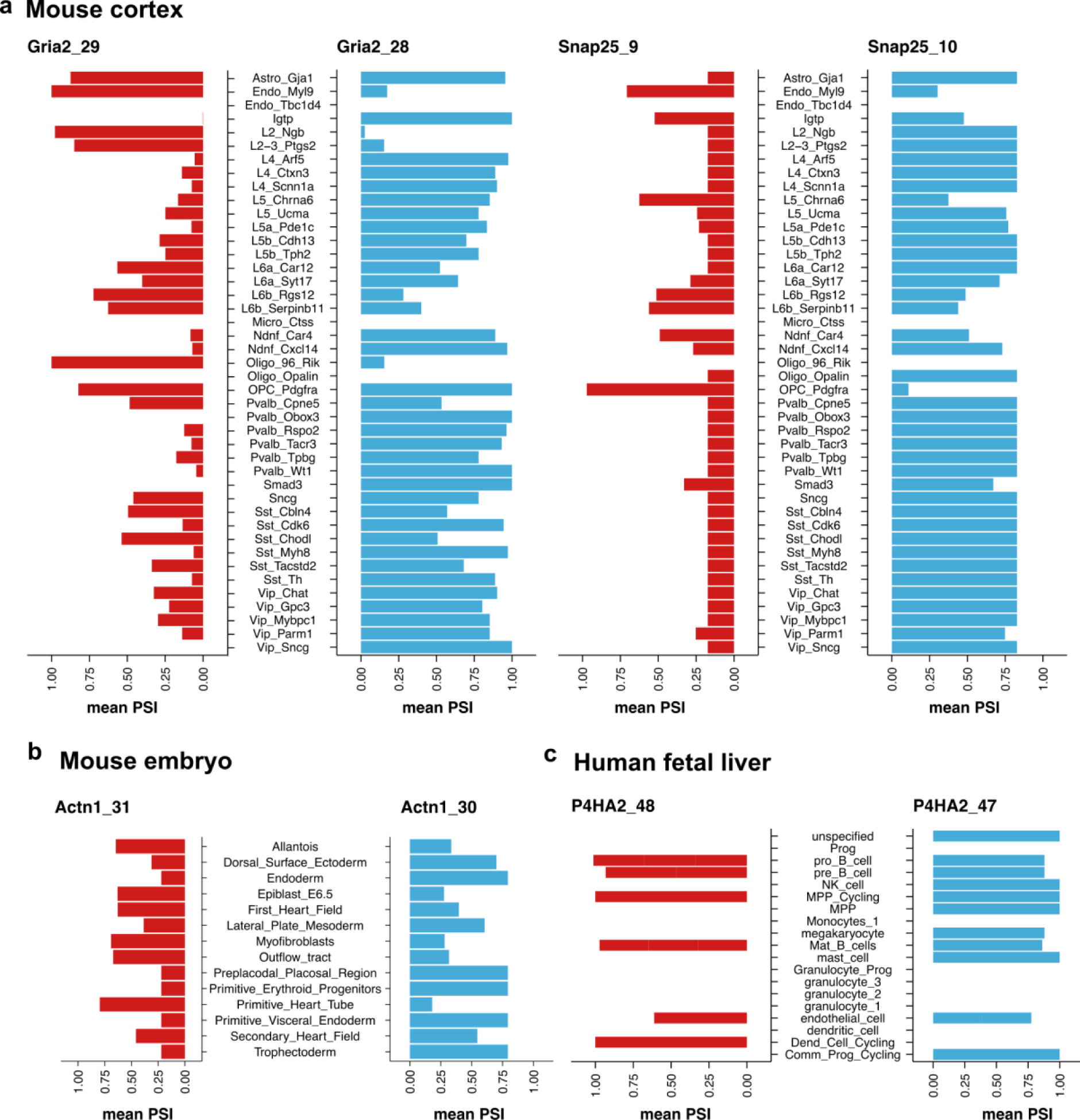
PSI values of detected high confidence adjacent mutually exclusive exons. The mean PSI across cell pools in each cell type in (a) *Gria2* (left) and *Snap25* (right) in the mouse cortex dataset, (b) *Actn1* in the mouse embryo dataset, and (c) *P4HA2* in the human fetal liver dataset. PSI, percent spliced-in.

Observing the cell type mean PSI of high-confidence MXEs, we found that the mutually exclusive pattern is not strictly followed in some cell types (**Fig 5**). For example, Sst_Cbln4 cells showed comparable PSI for nodes 28 and 29 in *Gria2*, whereas astrocytes included both exons. This is in parallel with previous observations of MXEs only being mutually exclusive in specific tissues but not in all tissues (Nilsen & Graveley, 2010; Pohl et al., 2013; Sammeth et al., 2008). For example, *TCL6* only shows exclusive patterns in specific tissues while on the basis of all known transcripts, the pattern is lost (Nilsen & Graveley, 2010; Pohl et al., 2013; Sammeth et al., 2008). Similarly, our results concurred that MXEs can show a mutually exclusive pattern only in some cell types.

### Identification of coordinately spliced exon blocks

By definition, the splicing nodes considered by scASfind only involve a single event, and consequently, the impact on the protein is likely to be limited. However, splicing events are often coordinated, resulting in multiple consecutive exons being simultaneously included or excluded, and we refer to such groups as node blocks. Coordinated events are more likely to have a more substantial impact on protein function as a larger proportion of coding sequences are affected, but when using a splicing node representation, they are difficult to detect as one must find a stretch of consecutive nodes. Given the large number of nodes across the transcriptome and the high noise level of splicing quantification, this search can be very time-consuming.

We used scASfind to detect node blocks, and for each gene, we first identify consecutive nodes of type ‘core exon’ with similar mean and standard deviation of their PSI values (the default is to require the absolute differences for both to be < 0.1 for all exons in the block). Next, the search is expanded to identify additional neighboring nodes to find cell type specific blocks. Events where a block of nodes is coordinately spliced-in (the ‘above’ events) and spliced-out (the ‘below’ events) are detected separately. To ensure high quality results, we only keep blocks composed of at least three nodes from different actual exons. Cell type specific node blocks that are detected in over half of the cell pools are reported as high confidence blocks.

Overall, we detected 263 node blocks with lengths ranging from 3 to 21 in the mouse cortex dataset, with 8 high-confidence ones (3-5 splicing nodes long). For the mouse embryo data, we found 306 blocks containing 3-26 nodes and 14 high-confidence ones with lengths ranging from 3 to 6. For the human fetal liver data, there were 526 node blocks ranging from 3 to 37 nodes, 19 of which were high confidence (with lengths from 3-9) (**Fig 6A-F**). For example, in the mouse cortex data, we found that node 5-7 in *Haus7* is significantly spliced-out in L2_Ngb and L5_Pde1c while it is spliced-in in *Sncg* (**Fig 6D**). Reassuringly, this is in line with two documented isoforms as shown in the Gencode annotation (**Extended data fig 1**). In the human fetal liver, we found a block of exons between chr2:95895399-95901206 in *ANKRD36C* that is spliced-in for dend_cell_cycling, also in concordance with known isoforms (**Extended data fig 2**).

**Figure 6.**
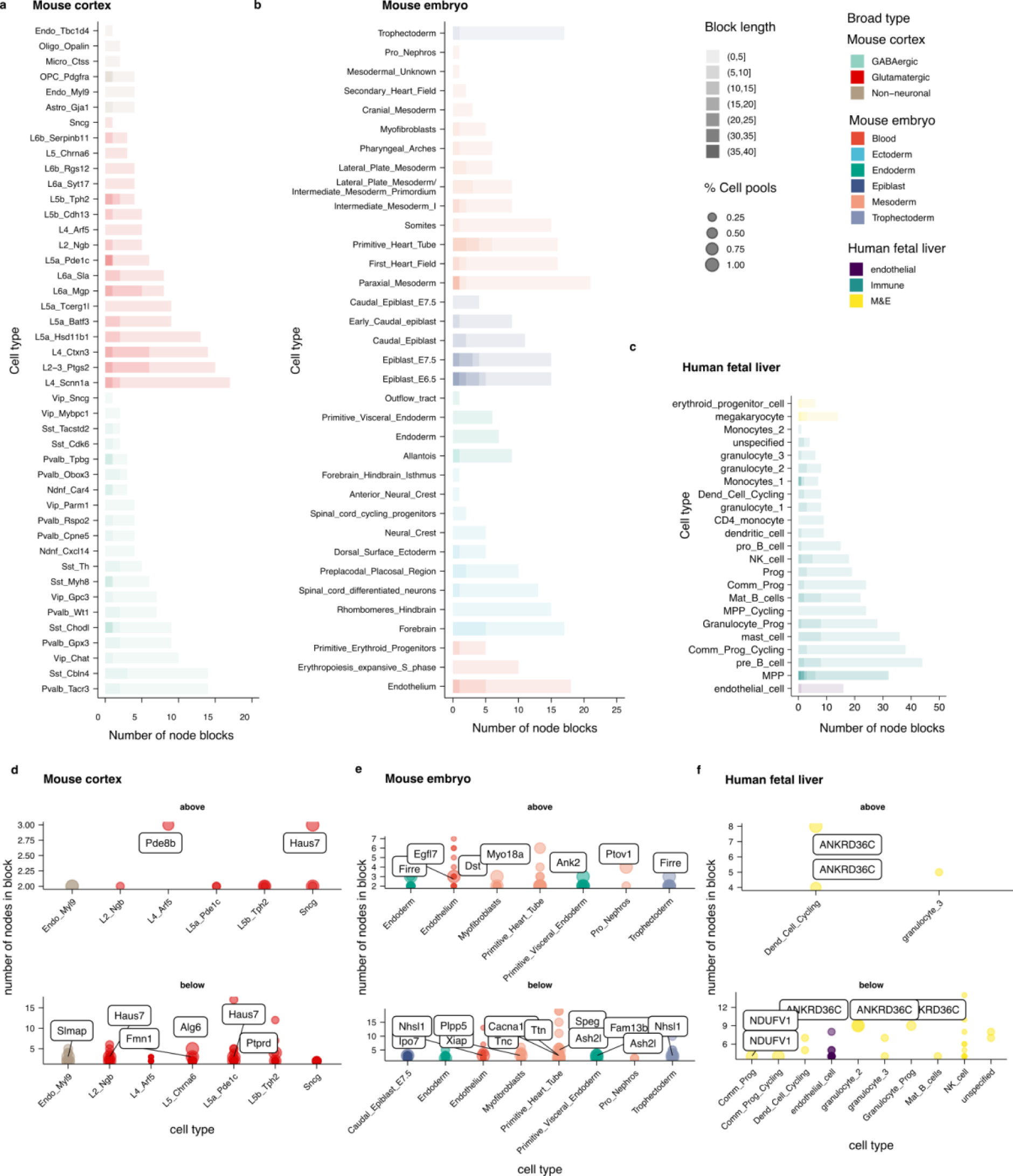
Cell type-specific coordinated splicing. Overall, (a) 263 node blocks were detected in the mouse cortex dataset, (b) 306 in the mouse embryo, and (c) 526 in the human fetal liver. The gene name of highly confident node blocks, which includes at least 3 nodes from different exons and were detected in >=50% cell pools are shown in (d-f) for each dataset. In d-f, ‘above’ means that the node block is significantly spliced-in in the respective cell type and ‘below’ means significantly splice-out of the node block.

We proceed to analyze whether the detected highly confident node blocks correspond to known isoforms, and what are their functional implications on protein domains. In the mouse embryo data, we detected known coordinated events in *Ttn* in primitive_heart_tube and first_heart_field. This corresponds to a regulated isoform switching event during heart development, upon which the stiffness of the protein changes (Lahmers et al., 2004; Opitz et al., 2004). Another key gene for heart development is *Dst*. Here we found a block including 5 exons and spanning 3 of the 7 major protein domains significantly spliced-in in myofibroblasts (**Fig 7A**). This event is in line with the documented muscle-specific Dst-b isoform (Leung et al., 2001). Mutation studies have shown that Dst-b is essential for strained muscle maintenance (Yoshioka et al., 2022). We have also found a node exclusion block in *Myo18A*, which is consistent with shorter annotated isoforms that are derived from an internal transcription start site (Guzik-Lendrum et al., 2013; Ouyang et al., 2021)(**Fig 7B**). One of these shorter isoforms, known as Myo18Aγ, lacks the PDZ-containing N-terminus but includes an alternative N-terminal extension (Horsthemke et al., 2019) and showed well-marked cell type specific profile associated to primitive_heart_tube. In general, the detected highly confident node blocks correspond well to known regulated isoforms. The analysis demonstrated the versatility of scASdfind to detect diverse isoform switching events.

**Figure 7.**
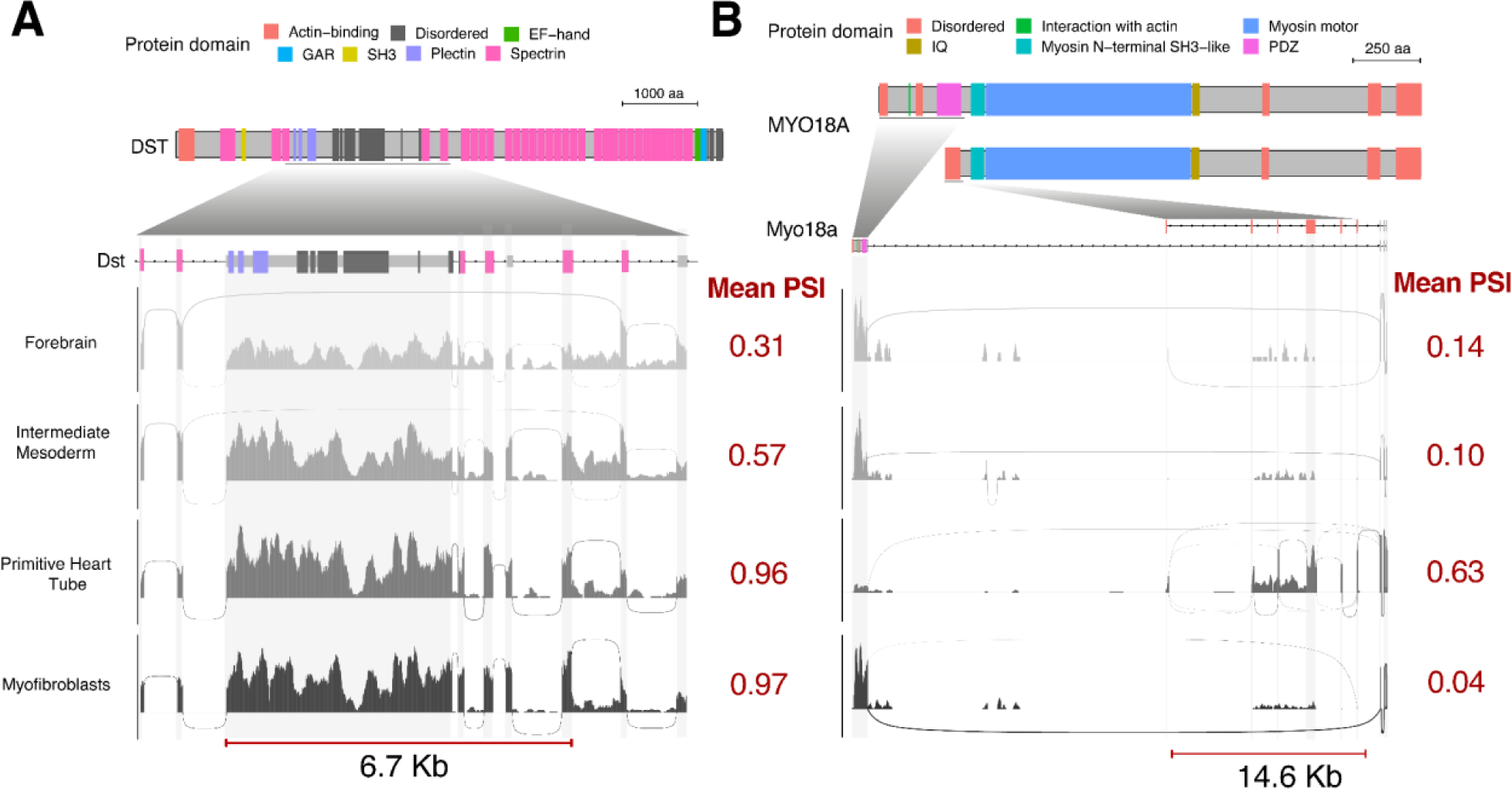
Coordinated inclusion of splicing nodes across mouse embryonic cell-types. Sashimi plots showing the read coverage and splice site usage across (a) Dst and (b) Myo18a transcripts. Top schematics show domains annotated for these proteins and coordinated splice nodes are highlighted with a red segment (bottom). Mean PSI values across all coordinated splice nodes are indicated in red. PSI, percent spliced-in.

Even though some blocks are detected in <50% of cell pools, they often match documented isoforms in the annotation. For example, in the human fetal liver, 20% of NK_cell cell pools did not include chr6:42632552-42655723 (14 splicing nodes) in the *UBR2* gene, in line with an isoform corresponding to early termination (**Extended data fig 3**). We also found that 25% of endothelial cells express a shorter isoform of *TBC1D19* (**Extended data fig 4**). Moreover, we have detected several high confidence blocks that suggest undocumented isoforms. For example, four exons between chr7:44864913-44865509 in *Ptov1* are shown to be coordinately included in all cell pools of Pro_Nephros in mouse embryos while there are no recorded isoforms of this gene in the Gencode annotation (**Fig 6F**, **Extended data fig 5, 6**). This could have relevance to human biology since *PTOV1* has been associated with prostate cancer, and the splicing event is likely to have a disruptive impact on one of the two major domains involved in the interaction with multiple other genes (Cánovas et al., 2017).

## Discussions

Splicing is a highly regulated process with a key role in cellular identity and function. Here we present scASfind, a toolkit for mining cell type-specific splicing patterns from large, single-cell, full-length transcriptomics datasets. Since many isoforms are highly overlapping, analysis of splicing patterns is challenging with short reads. With single cell data it is even more difficult due to the much larger number of isoforms compared to genes, and the much sparser coverage of each splicing node. To overcome these challenges, we utilized pooling and compression to build an index which can support queries of what cell types are enriched for a set of splicing events. We demonstrate that an index for thousands of cells can be created in 20-30 minutes, resulting in compression by 2-3 orders of magnitude, while at the same time speeding up queries hundreds of folds. Building on this data structure we provide functions for discovering cell type-specific splicing events such as finding marker nodes, mutually exclusive exons, or coordinated node blocks. Using mouse cortex, mouse embryonic development, and human fetal liver datasets, we demonstrated scASfind’s utility for carrying out tasks that would have been prohibitive without the tool.

Isoform quantification is a very challenging problem with short-read data, but quantification of splicing events is a more tractable challenge. In combination with the fact that the most widely used single-cell platforms only allow for the 3’ or 5’ end of the transcript to be sequenced, this means that alternative splicing remains a poorly explored aspect at the single-cell level. This is likely to change in the near future due to high throughput full-length protocols such as VASA-seq (Salmen et al., 2022). Moreover, several studies have utilized long reads technologies for single-cell studies (Lebrigand et al., 2023; Philpott et al., 2021; Tian et al., 2021), allowing an entire transcript to be captured by a single read. We believe that these advances will allow single-cell studies to quantify alternative splicing events, but for this to become feasible novel computational methods are required. Given its efficient memory usage, low run times and convenient search functionality, we believe that scASfind will be a valuable tool for researchers to decipher cell type-specific splicing using scRNA-seq data.

## Methods

### AS quantification across cell-types

To quantify AS events across cell-types we configured and ran MicroExonator’s single-cell module, as described in (Parada & Hemberg, 2022). As part of this workflow, MicroExonator quantifies AS events using Whippet (Sterne-Weiler et al., 2018) across pseudobulbs derived from annotated cell clusters. Using this protocol, we processed mouse scRNA-seq data from brain visual cortex (Tasic et al., 2016) and whole embryos (Salmen et al., 2022), as well as scRNA data derived from human immunophenotypic blood cells from fetal liver and bone marrow (Ranzoni et al., 2021). We used genome assembly mm10 and GENCODE transcript annotation v16 to process mouse scRNA-seq data. For human scRNA-seq analyses, we used genome assembly hg38 and GENCODE transcript annotation v19.

### Filter for confidently quantified events

Before encoding the PSI data, we first filter for confidently quantified events. The sparsity of scRNA-seq data often results in an insufficient number of reads spanning splicing junctions that can be used to calculate node PSI values. By default, we require at least 10 reads available for PSI quantification. This gives roughly a confidence interval of PSI < 0.5 from Whippet.

### Create a scASfind index

scASfind builds four types of data structures from the input data, and together they form a queryable index.

1. The splicing node x cell pool differential PSI matrices For each node, we first calculate the mean PSI across all cell pools, then calculate the difference from the mean for all PSI values. A 0.2 deviation was used as the default threshold to select sufficiently deviated events. Secondly, we separate nodes with differential inclusion (the ‘above’ events) and those with differential exclusion (the ‘below’ events). Both metrics are then multiplied by 100 so that the value is in the range of [0,100] for the compression. The splicing node x cell pool differential PSI (ΔPSI) matrices are independently compressed using the strategy in scfind (Lee et al., 2021). The compression is a two-step process: 1) storing the positions of non-zero values are compressed by Elias-Fano encoding, and 2) the actual differential PSI value is represented as quantiles of a log-normal distribution (extended data figure 7). The mean and variance of the log-normal distribution, along with quantiles of the original ΔPSI value are stored. The first step is lossless while the second step is lossy. The approximation of actual ΔPSI in the index makes it possible to retrieve the approximate PSI value when giving the user the ability to tune the size of the storage based on the number of bits used for the quantization (default 2 bits).
2. The mean and standard deviation PSI values per node We store the mean and standard deviation for each dataset and node for retrieval of raw PSI values, and for speeding up searches of MXEs and node blocks based on expected patterns in mean and standard deviation.
3. The mask for NA values from PSI quantification Since we only encode differential PSI values, cell pools with PSI values close to the dataset-wise mean are excluded. However, cell pools where the PSI value is unquantified (NA) or below the required number of reads for confident quantification are also excluded. Distinguishing these two circumstances is required to enable retrieval of raw PSI values from the index. For this purpose, we use a binary mask matrix. In this matrix, 1 indicates the cell pools with PSI value equal to the dataset-wise mean and 0 indicates unquantified. Typically, unquantified events are more frequent than equal mean events, resulting in a sparse matrix.
4. The annotation of nodes We use ENSEMBL (Cunningham et al., 2022) via the R package biomaRt (V2.46.3) to obtain annotations of all nodes included in the index to enable quick interpretation of results.

In addition, metadata for each cell, providing information about its annotated cell type or state is required. The buildAltSpliceIndex function in scASfind takes the cell pool-by-splicing node differential PSI matrices and a table with the cell type annotation to build an index object. The ‘above’ and ‘below’ index objects are stored as two datasets and the three other metrics are stored in the metadata slot in the scASfind object.

### Benchmark of file size, index build time and node search time

We benchmark the efficiency of the scASfind index, compared with a basic approach utilizing only R and Seurat functions. For file size, we take the sum of on-disk space taken by the raw PSI matrix (as .tsv files) and the three metadata objects (as .rds files) as “raw PSI”, and the complete scASfind index (as .rds object), including the same three metadata objects stored in the metadata slot as”‘scASfind index”. All file sizes are obtained with the “file.info” function in R. For index build time, we run the scASfind build index script on a high-performance computing cluster (Rocky Linux 8.5) for the three datasets with 4 cores and maximum 2GB memory, 10 processes and 200 threads. The time to build the index naturally depends on the computational resources available. For differential events search time, we randomly select 5, 10, 50, 100 and 200 splicing nodes, and search for cell pools with an above-mean PSI of any of the nodes using either the naive approach or the scASfind index. The elapsed times were measured in 30 repetitions per query length.

### Cell type marker nodes search

We use a precision-recall framework to search for nodes that are specific to different cell types. For each node, we count the number of cell pools with the relative inclusion/exclusion of this node in a cell type of interest compared to all other cell types. We use precision, recall and F1 scores to evaluate how well the node distinguishes the cell type of interest from all other cell types. A true positive (TP) is when a node is included or excluded in a cell pool from the cell type of interest for spliced-in and spliced-out events, respectively. False positives (FP) are when the same node is included/excluded in cell pools of other cell types, and false negatives (FN) are cell pools from the same cell type in which the node is not detected as included or excluded. The precision score is calculated by:

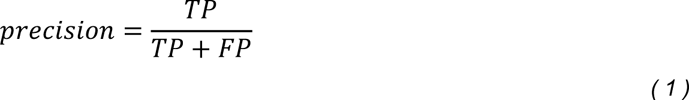

The recall score is calculated by:

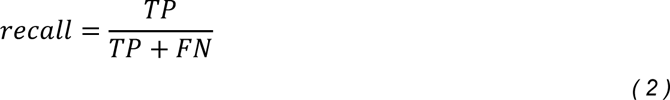

The F1 score is the harmonic mean of precision and recall score:

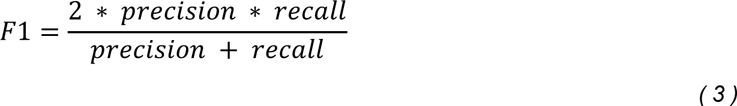

By default, we use F1 scores as a balanced metric to rank all nodes for each cell type to indicate the best marker nodes for either inclusion or exclusion events.

### Comparing PSI and gene expression in splicing marker nodes

We calculate the top 20 gene expression markers (ranked by F1 score) using scfind in the mouse cortex data. For the genes containing splicing marker nodes, we used the R package Seurat (V4.1.0) to obtain the scaled expression values. Then we used ggplot2 (V3.3.3), viridislite (V0.4.0) and cowplot (V1.1.1) to create the violin plot of PSI and scaled expression values.

### Detect cell type-specific mutually exclusive exons

Detection of cell type-specific MXEs is based on a hypergeometric test with the ‘hyperQueryCellTypes’ function in scASfind. The hypergeometric distribution models the probability of *k* success in *n* draws without replacement, from a finite population with *N* subjects and *K* of them contains the pattern. In our case, *k* is the number of cell pools in a cell type which have the splicing pattern, *n* is the number of cell pools in that cell type, *K* is the total number of cell pools in which the splicing pattern is detected and *N* is the total number of cell pools. A pattern with a hypergeometric test P-value <= 0.05 in a cell type is considered significant.

We use a mutually exclusive combination of splicing nodes (include one and exclude the other) as the pattern in the hypergeometric test to detect MXEs. The query is performed exhaustively for all pairs of exons in the dataset. To reduce the search space, we first filter all the possible node pairs by 1) having a mean PSI sum of 1±0.1 and 2) having a <0.1 difference in the PSI standard deviation. Then, we query for significant cell types for the potential pairs of MXEs. Pairs with at least one cell type significant in one of the two possible patterns are kept as candidates.

Further filters are applied for candidate MXE pairs. First, we require the MXE pattern to be found in >= 50% of the pools in the significant cell type. Then, we require the difference of absolute PSI value in the pair to be >= 0.5 to be considered high confidence.

### Detecting cell type-specific coordinately spliced-in exons

We scan all genes from the 5’ of all annotated ‘core exon’ nodes in the scASfind index to find coordinately spliced-in exons. We extend an exon block by requiring the next exon to have at most ± 0.1 difference with both the mean and standard deviation of the previous block of exons. If this criterion is not fulfilled, we initiate a new block, and the previously constructed block is tested for cell type specificity using the hypergeometric test as described previously. Blocks significant in at least one cell type are kept. Finally, we use a node-to-exon mapping table from Whippet (Sterne-Weiler et al., 2018) to combine nodes belonging to the same actual exon. If the resulting block contains at least 3 exons, we propose it as a potential coordinated sliced-in exon block. We also require the block to be found in >50% of the pools in the significant cell type for it to be highly confident.

## Data and code availability

We provide scASfind freely available via zenodo (DOI: 10.5281/zenodo.8241682) or GitHub (https://github.com/hemberg-lab/scASfind) under an MIT license. MicroExonator is freely available via https://github.com/hemberg-lab/MicroExonator.

## Acknowledgements

We would like to thank Jae-Won Cho and Ruben Chazarra-Gil for feedback on the manuscript and Nikolaos Patikas for help in testing the software. YS was funded by the EMBL international PhD program, GP and JTHL were funded by the Wellcome Trust, and MH was funded by the Wellcome Trust and the Evergrande Center.

## Author Contributions

GP, JTHL and MH conceived the project. YS wrote the code and analyzed the data with assistance from GP and JTHL. YS and MH wrote the manuscript with input from GP and JTHL. MH supervised the research.

## Extended Data

**Extended data figure 1:**
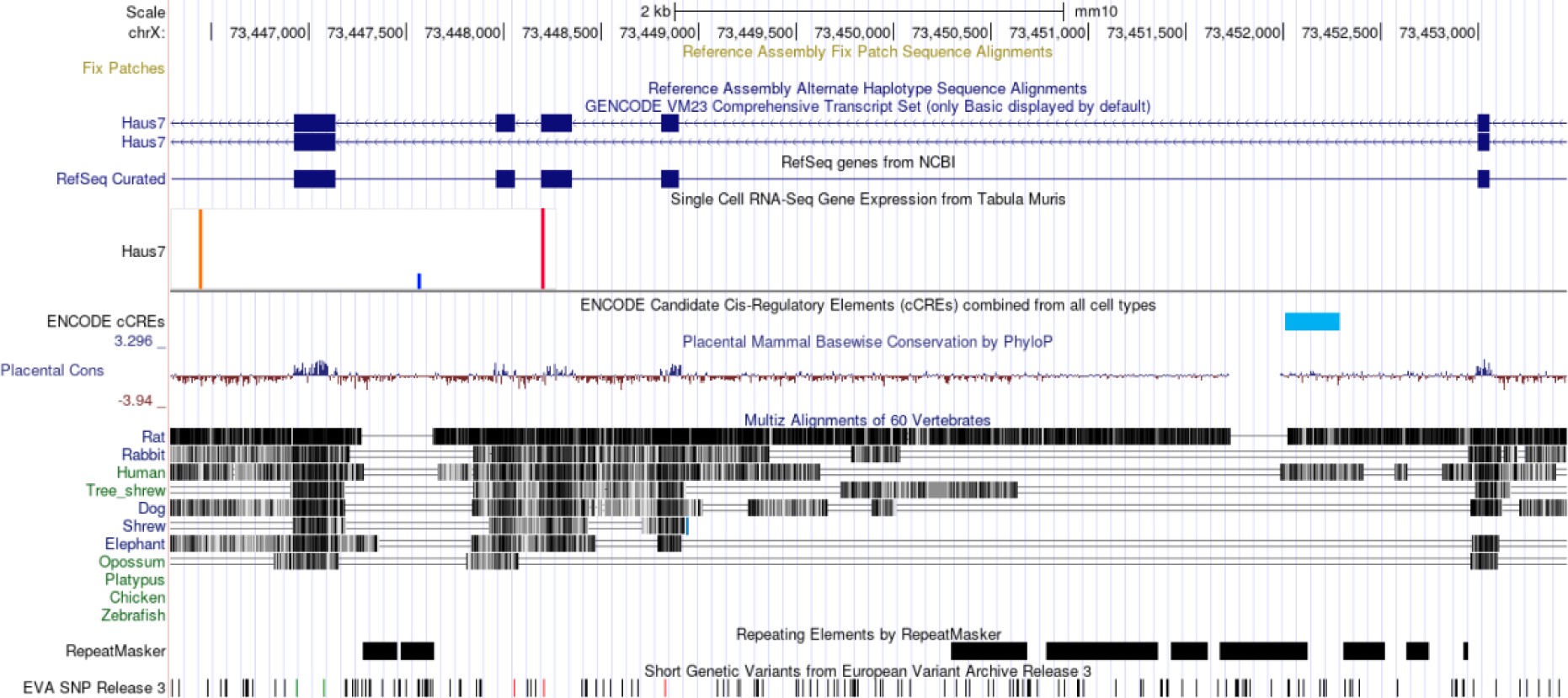
Reference assembly of *Haus7* in the mouse genome from the UCSC genome browser. Showing the two alternative isoforms with node blocks corresponding to scASfind results.

**Extended data figure 2:**
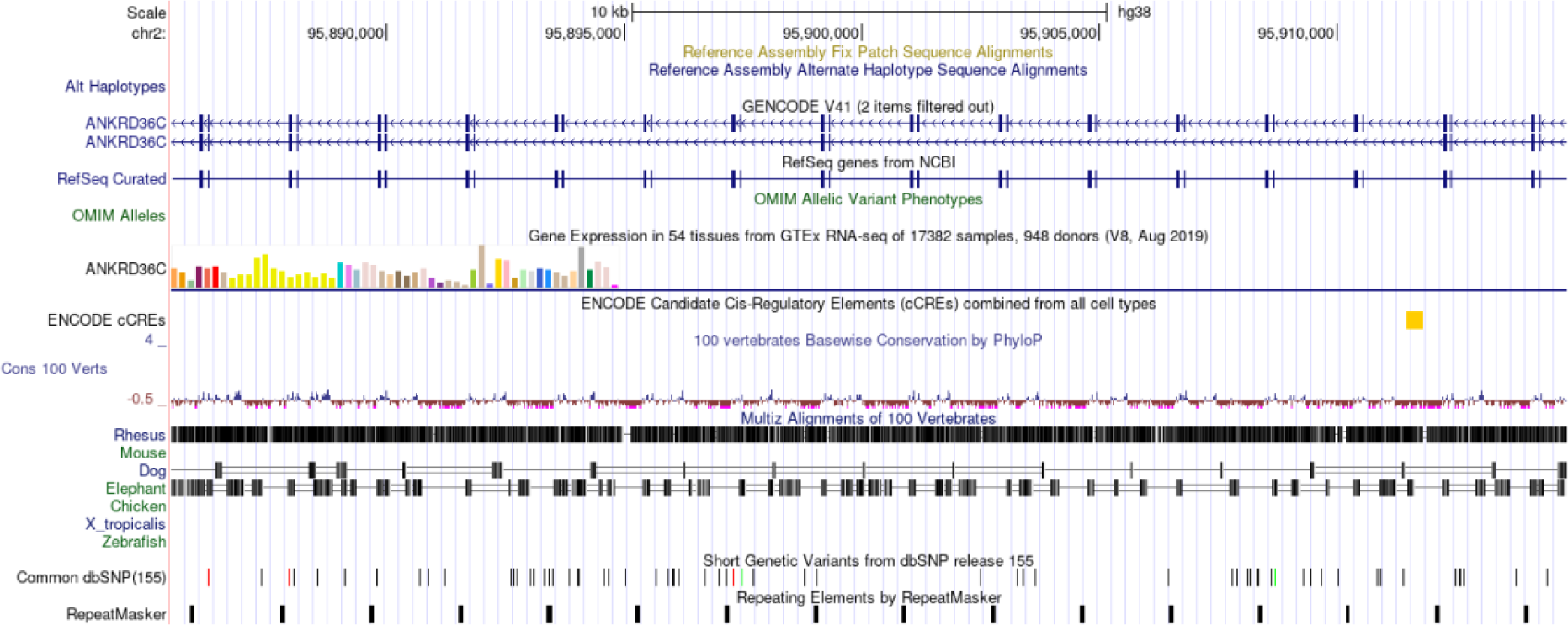
Reference assembly of *ANKRD36C* in the human genome from the UCSC genome browser. Showing the two alternative isoforms with node blocks corresponding to scASfind results.

**Extended data figure 3:**
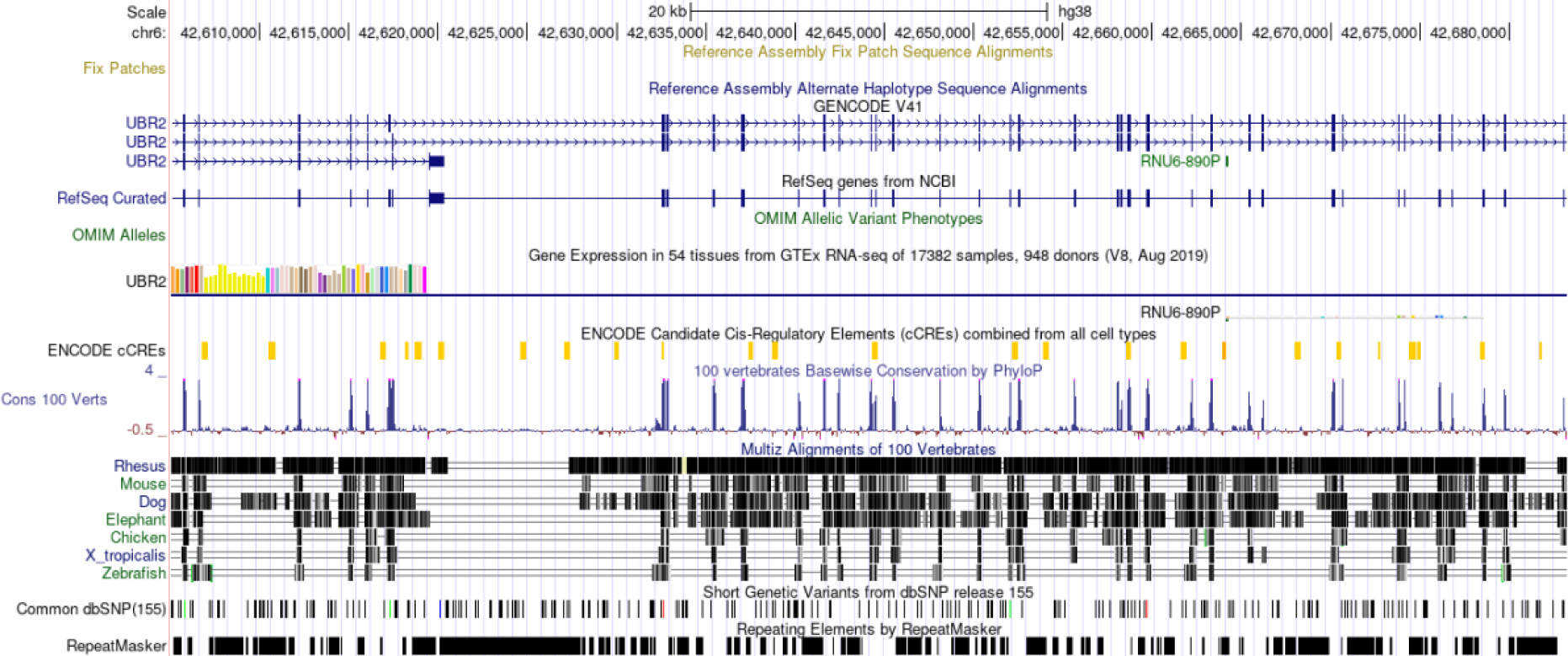
Reference assembly of *UBR2* in the human genome from the UCSC genome browser. Showing the three alternative isoforms of which one results in early termination, in line with scASfind results.

**Extended data figure 4:**
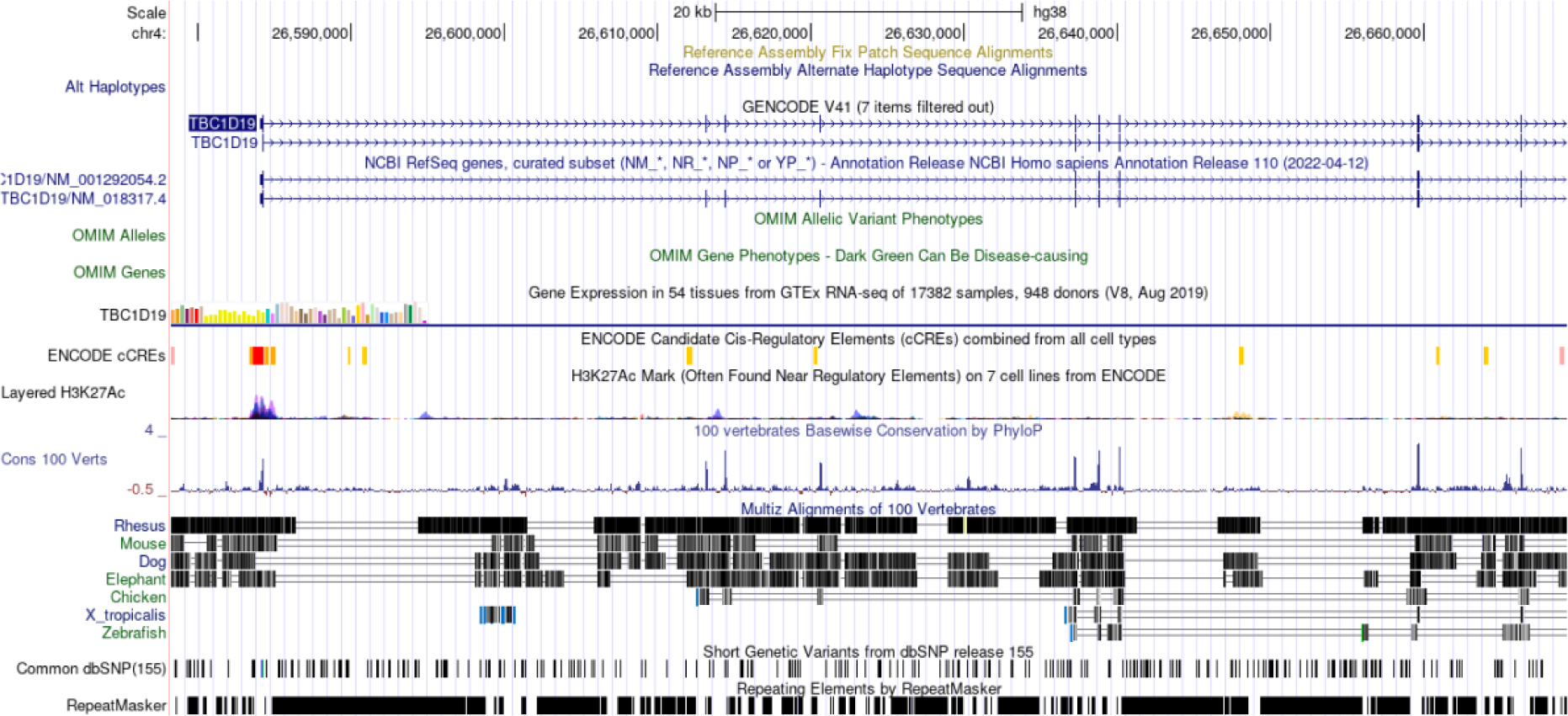
Reference assembly of *TBC1D19* in the human genome from the UCSC genome browser. Showing the two alternative isoforms with node blocks corresponding to scASfind results.

**Extended data figure 5:**
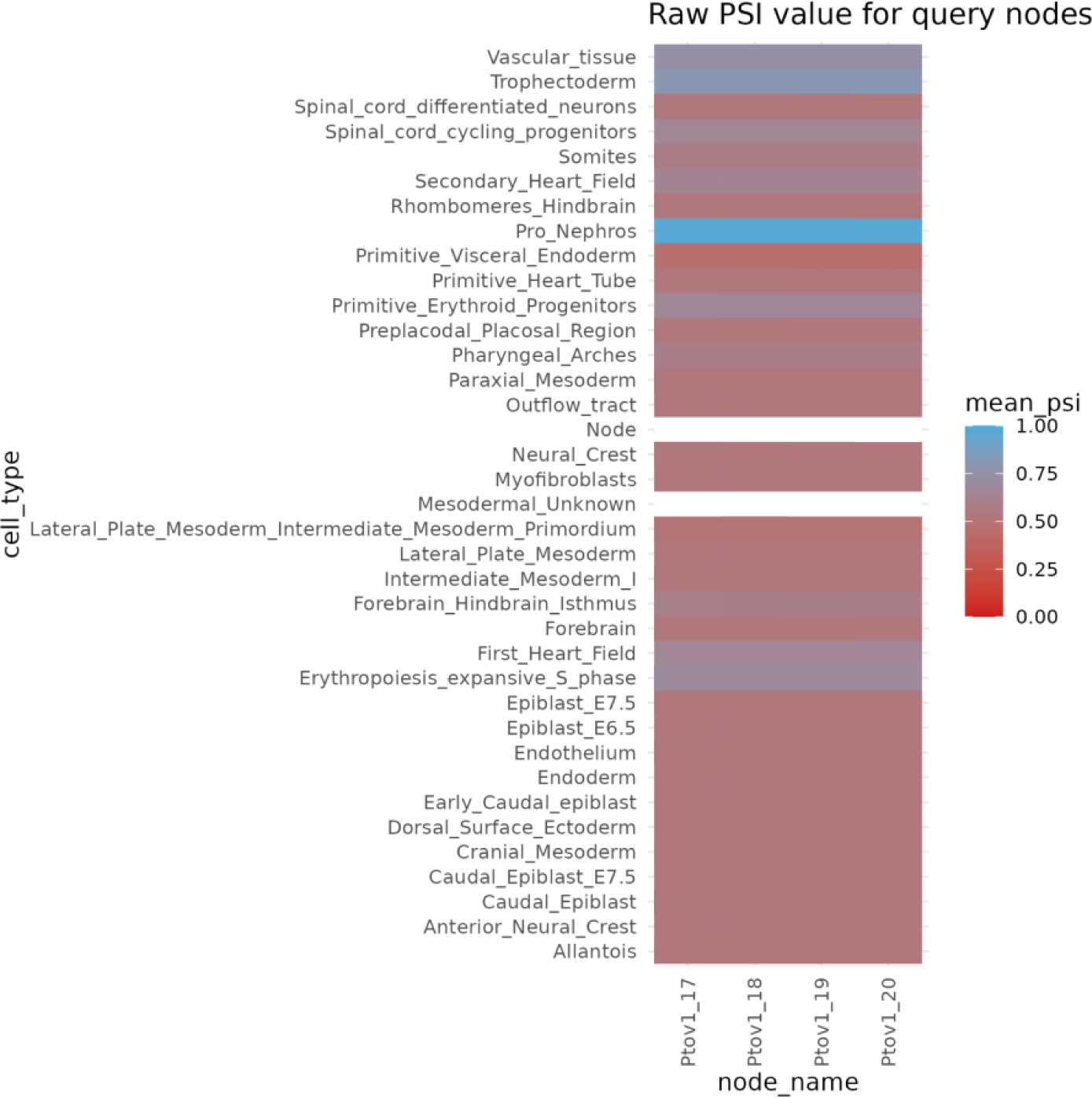
Raw PSI of node block in *Ptov1*. Showing inclusion of node *Ptov1_17*-*Ptov1_20* in Pro_Nephros in the mouse embryo data.

**Extended data figure 6:**
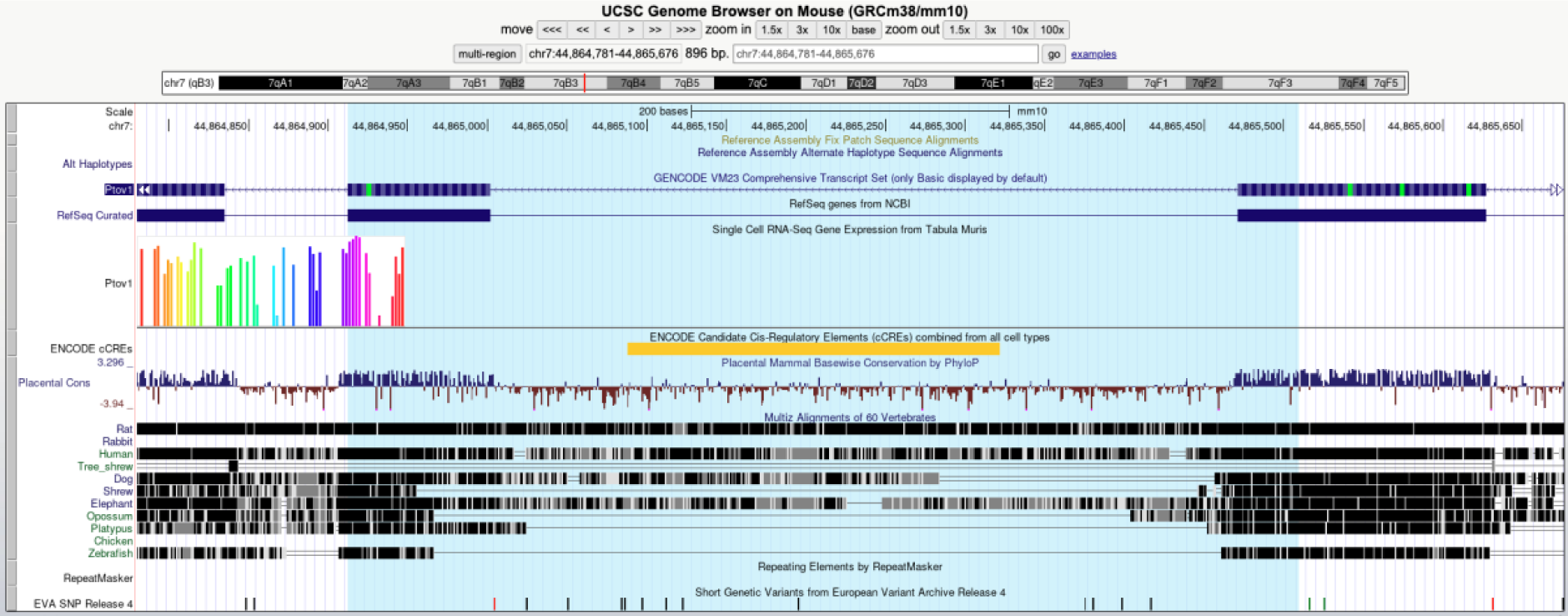
Ptov1 gene in mouse genome from the UCSC genome browser. Blue highlighted region is the detected node block by scASfind that has no documented isoform in the mm10 reference genome.

**Extended data figure 7:**
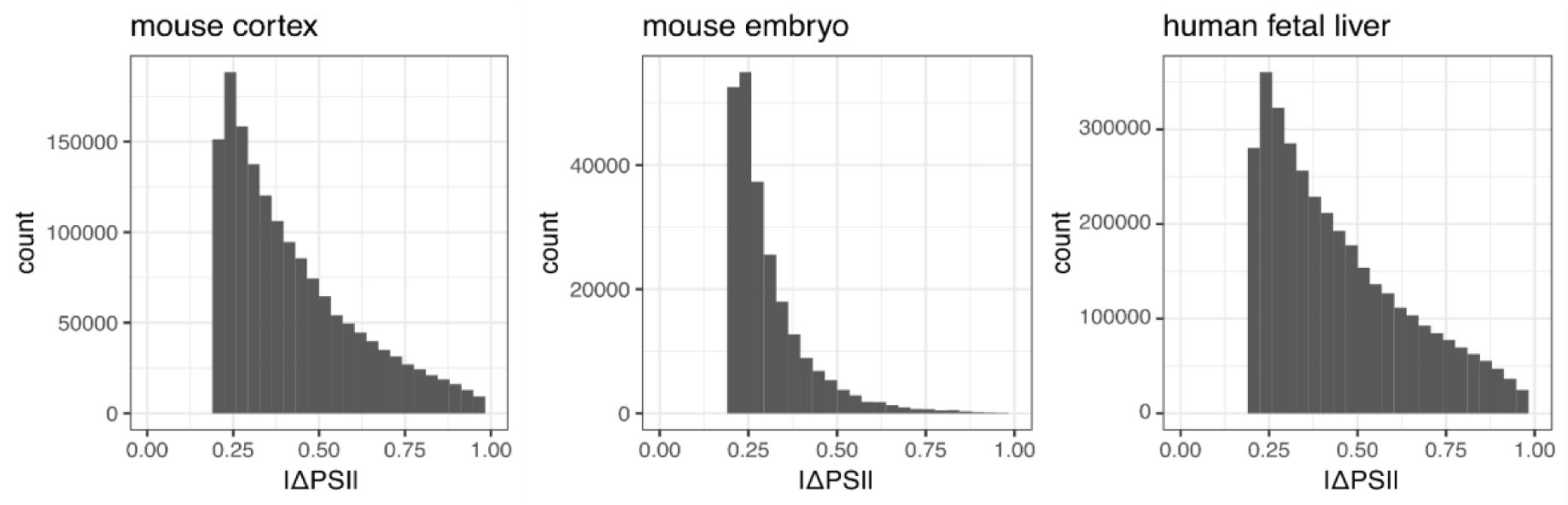
Histogram of PSI deviation from mean in the three datasets in all genes and all cell pools. Only showing encoded events which requires |ΔPSI| > 0.2. The histograms suggest that ΔPSI can be approximated by a lognormal distribution.

**Extended Data Table 1:** datasets used in this study, on-disk size and the time of building scASfind index. We pooled scRNA-seq reads from 5 cells, 200 cells and 10 cells to build cell pools for PSI quantification of the mouse cortex, mouse embryo and human fetal liver data, respectively. Splicing nodes which had a PSI deviating from the dataset mean in at least one cell pool were encoded. The size of raw data includes the raw PSI matrix and relevant metadata objects, while the size of the scASfind index object involves the compressed PSI index and the same metadata objects (see Methods). Both sizes are on-disk file sizes. The time to build the scASfind index is calculated by running the index- building script in scASfind on an HPC cluster.

| Dataset | Number of single cells | Number of cell pools | Number of cell types | Number of cell pools | Number of encoded splicing nodes | Size of raw data (MB) | Size of scASfind index object (MB) | Time to build the scASfind index (seconds) |
| --- | --- | --- | --- | --- | --- | --- | --- | --- |
| Mouse cortex (Tasic et al., 2016) | 1,654 | 339 | 49 | 339 | 90,834 | 14,142.8 | 55.5 | 1,185 |
| Mouse embryo (Salmen et al., 2022) | 33,662 | 191 | 37 | 191 | 75,837 | 27,484.3 | 21.9 | 1,366 |
| Human fetal liver (Ranzoni et al., 2021) | 4,503 | 451 | 23 | 451 | 199,361 | 29,374.5 | 101.4 | 1,864 |

